# Borderline personality disorder with cocaine dependence: impulsivity, emotional dysregulation and amygdala functional connectivity

**DOI:** 10.1101/268169

**Authors:** Thania Balducci, Jorge J Gonzalez-Olvera, Diego Angeles-Valdez, Isabel Espinoza-Luna, Eduardo A Garza-Villarreal

## Abstract

Objective: Borderline personality disorder (BPD) is present in 19% of cocaine dependence (CD) cases; however, this dual pathology (DP) is poorly understood. We assessed impulsivity, emotional dysregulation (ED) and amygdala functional connectivity in this DP.

Methods. We recruited 69 participants divided into 4 groups: DP (n = 20), CD without BPD (n = 19), BPD without CD (n = 10) and healthy controls (HC, n = 20). We used self-reported instruments to measure impulsivity and ED. We acquired resting state fMRI and performed seed-based analyses of functional connectivity (FC) of bilateral amygdalas.

Results. BPD and CD factors had opposing effects in impulsivity and ED, as well as on FC between left amygdala and medial prefrontal cortex. For the FC between right amygdala and left insula, the effect of having both disorders was additive, reducing FC strength. Significant FC clusters were correlated with impulsivity and ED.

Conclusions. In this preliminary study, we found that clinical scores of DP patients were closer to those of BPD without CD than to those of CD without BPD, while amygdala to medial prefrontal cortex FC patterns in DP patients were closer to HC than expected.

## Significant outcomes

- Patients with the dual pathology of borderline personality disorder and cocaine dependence present unique patterns of amygdala functional connectivity compared to those with only one of the disorders.
- The counteracting effects of borderline personality disorder and cocaine dependence on impulsivity, emotion dysregulation and amygdala to medial prefrontal cortex functional connectivity in the dual pathology patients raises the question of cocaine consumption as self-medication in patients with borderline personality disorder.
- Even when cocaine consumption could be a way to regulate amygdala to medial prefrontal cortex functional connectivity in patients with borderline personality disorder, other brain connections might deteriorate in these patients, for example the amygdala to insula functional connectivity.

## Limitations

- Small overall sample size and unequal groups
- Convenience sample of patients seeking care at specialized psychiatric and treatment units
- Reliance on self-report clinical measures and brain activity only assessed during resting state

## Introduction

When studying substances use disorders, psychiatric comorbidities are more the rule than the exception, being present in 85 to 95% of cases (1); the presence of such comorbidity is defined as dual pathology (DP) (2). Borderline personality disorder (BDP) with cocaine dependence (CD) as a DP has been rarely studied, even though 19% of patients with CD present with comorbid BPD (3). BPD is characterized by a pattern of instability of affect, self-image and interpersonal relationships, accompanied by impulsivity (4). Studies have found that CD is a consistent predictor for BPD with an odds ratio = 2.06, even higher than the risks generated by alcohol or opiate use disorder, substances that have been more widely studied in relation to BPD (5, 6). Cocaine is a stimulant amine and it is the second most used illicit drug in Mexico, Central America, Western Europe and South Africa (7). It has been estimated that 5-6% of those who consume cocaine develop dependence within the first year of use (8). This dependence is characterized by substance misuse with tolerance, abstinence syndrome, difficulties controlling consumption and clinical impairment or distress (4).

Women with this DP show higher sexual risk behaviors compared to women with CD only and to women with BPD and other substance use disorders (e.g. alcohol) (9). Men with this DP show a greater attention bias to cocaine-related visual stimuli under emotional stress when compared to men with CD only (10). As impulsivity and emotional dysregulation (ED) are present in both conditions separately, these traits have been suggested as possible etiological factors of vulnerability to develop this DP and to account for the between-group differences noted (9–11).

Impulsivity can be defined as the tendency towards rapid unplanned reactions to internal or external stimuli without considering the consequences (12). Impulsivity increases risk for stimulant use disorder (13) and is a predictor for lifetime cocaine use (14). In BPD, impulsivity was the strongest predictor for borderline psychopathology over a seven year follow-up (15). ED is the difficulty to control and modulate one’s affective state, such that emotions escape rational control and judgment (16). It is a core dimension of BPD and is present during drug abstinence in CD patients (17).

Brain imaging studies have found several structures with altered functional connectivity (FC) in both disorders separately. In BPD, during an emotional processing task, amygdala activity was increased when exposed to fearful faces, while anterior cingulate cortex (ACC) activity was decreased. The opposite activation pattern was shown for angry faces (18). Another study found hyperreactivity of amygdala, ACC and insula for negative and neutral pictures (19). In resting state, studies have found higher FC among amygdala, insula and orbitofrontal cortex, and lower FC between ACC and posterior cingulate cortex (PCC). In CD, lower FC has been found between amygdala and medial prefrontal cortex (mPFC) and between ACC and posterior insula (20). Default mode network (DMN) connectivity seems to be also disrupted in both disorders, being higher between mPFC and precuneus/PCC in BPD (21), while greater FC density in ventromedial prefrontal cortex, precuneus and PCC has been found in CD (22). Amygdala, mPFC, ACC, PCC, precuneus and insula are all involved in emotional regulation (23–27). However, the roles of impulsivity and ED in this DP are unknown and there are no published FC studies addressing this DP.

### Aims of the study

To investigate impulsivity, emotional dysregulation and functional connectivity in dual pathology (borderline personality disorder with cocaine dependence) compared to the single pathologies and to healthy controls. We hypothesized additive effects of the two disorders in the clinical domain, and negative interactions in areas related to functional connectivity dysfunction, that is, we expected opposing effects of each disorder on amygdala functional connectivity.

## Material and methods

### Participants

The sample consisted of 69 participants divided into four groups: 20 with dual pathology of cocaine dependence and borderline personality disorder (BPD+CD+), 19 with cocaine dependence without borderline personality disorder (BPD-CD+), 10 with borderline personality disorder without cocaine dependence (BPD+CD-) and 20 controls without psychopathology (BPD-CD-). Demographic characteristics are summarized in Table 1. Participants were recruited from the out-patient Addiction Clinic and the Borderline Personality Disorder Clinic at the Instituto Nacional de Psiquiatría “Ramón de la Fuente Muñiz” and from the Xochimilco Toxicological Medical Unit (treatment clinic) in Mexico City. The patients were a subsample from an ongoing cocaine addiction study (28). The BPD-CD- group was recruited via flyer advertisements and word of mouth. The groups were matched for sex, age, handedness and economic status. All subjects provided written informed consent. The study follows the guidelines outlined in the Declaration of Helsinki and was approved by the Ethics Committee of the Instituto Nacional de Psiquiatría “Ramón de la Fuente Muñiz”.

**Table 1.** Demographic characteristics of the study participants

| | BPD+CD+<br>(n = 20) | BPD-CD+<br>(n = 19) | BPD+CD-<br>(n = 10) | BPD-CD-<br>(n = 20) | $F/\chi^2$<br>value | *p* value |
| --- | --- | --- | --- | --- | --- | --- |
| Age, years (SD) | 31 (7) | 31 (6) | 31 (13) | 32 (8) | 0.173 | 0.914 |
| Gender: male, n (%) | 15 (75) | 18 (94) | 5 (50) | 18 (90) | 10.16 | < 0.05 |
| Education, median | High school | Junior high school | Technical degree | Technical degree | 11.98 | < 0.01 |
| Economic status*, median | D+ | C | C/C+ | C | 4.071 | 0.254 |
| Employment, n (%) |  |  |  |  |  |  |
| Full-time | 7 (35.0) | 8 (42.1) | 2 (20.0) | 10 (50.0) | 22.27 | 0.220 |
| Half-time, formal | 2 (10.0) | 4 (21.1) | 2 (20.0) | 2 (10.0) |  |  |
| Half-time, informal | 5 (25.0) | 5 (26.3) | 2 (20.0) | 2 (10.0) |  |  |
| Student | 1 (5.0) | 2 (10.5) | 4 (40.0) | 4 (20.0) |  |  |
| Housekeeper | - | - | - | 1 (5.0) |  |  |
| Home | 2 (10.0) | - | - | - |  |  |
| Unemployed | 3 (15.0) | - | - | 1 (5.0) |  |  |
| Marital status, n (%) |  |  |  |  |  |  |
| Unmarried | 7 (35.0) | 8 (42.1) | 6 (60.0) | 10 (50.0) | 3.331 | 0.766 |
| With partner | 6 (30.0) | 6 (31.6) | 2 (20.0) | 7 (35.0) |  |  |
| Divorced/separated | 7 (35.0) | 5 (26.3) | 2 (20.0) | 3 (15.0) |  |  |
| Laterality: right-handed, n (%) | 19 (95.0) | 14 (73.7) | 10 (100) | 17 (85.0) | 6.392 | 0.381 |

\*The instrument used was the AMAI rule 8x7 created for Mexican homes, where A/B is the highest economic status category and E is the lowest.
BPD: borderline personality disorder, CD: cocaine dependence

For BPD screening, we used the self-report version of the Structured Clinical Interview for the Diagnostic and Statistical Manual of Mental Disorders 4th edition Axis II and the diagnosis was made with the Diagnostic Interview for Borderline Revised administered by a psychiatrist trained on personality disorders. Cocaine dependence was diagnosed using the MINI International Neuropsychiatric Interview Spanish version which was administered by two attending psychiatrists and two third year psychiatry residents who were supervised by the psychiatrists.

The MINI International Neuropsychiatric Interview was used also to diagnose psychiatric comorbidity. Subjects with bipolar, psychotic, obsessive-compulsive and eating disorders were excluded. For BPD+CD- and BPD-CD- groups, the presence of any substance abuse or dependence except nicotine was an exclusion criterion. BPD+CD+ and BPD-CD+ groups could have another substance use disorder if cocaine was the primary substance. Cocaine consumption had to be active or with abstinence less than 60 days prior to the scan, with frequency of use of at least three days per week and no more than 60 continued days of abstinence during the last 12 months. Additional exclusion criteria for all groups were somatic diseases, including neurological disorders, severe suicidal risk, history of head trauma with loss of consciousness, pregnancy, obesity and noncompliance with magnetic resonance imaging safety standards. BDP-CD- presenting any psychiatric or somatic disorder were excluded.

### Clinical measures

Self-reported impulsivity was evaluated with the Barratt Impulsiveness Scale (BIS-11) which has three subscales: non-planning impulsiveness, which involves a lack of forethought; cognitive impulsivity, which involves making quick decisions; and motor impulsivity, which involves acting without thinking (29). ED was assessed with the Difficulties in Emotion Regulation Scale (DERS) validated in Mexico (30), which unlike the original version, has 24 items and five subscales: non-acceptance of emotional responses, difficulty engaging in goal-directed behavior, lack of emotional awareness and lack of emotional clarity. Severity of CD was assessed with the Addiction Severity Index. For the CD+ groups, craving at the time of the magnetic resonance imaging acquisition was evaluated with the Cocaine Craving Questionnaire-Now. The severity of BPD was assessed using the Clinical Global Impression Scale for BPD.

### Magnetic resonance imaging acquisition

Imaging data were obtained at a 3.0 T Philips Ingenia magnetic resonance imaging scanner with a 32-channel phased array head coil. For the resting state fMRI, participants were instructed to remain quiet, relax and fixate on a cross. T2*-weighted echo planar images were acquired during 10 minutes (300 axial slices, repetition time = 2000 ms, echo time = 30 ms, flip angle = 75°, field of view = 240 mm, slice thickness = 3.0 mm, acquisition matrix = 80 × 80 and voxel size = 3.0 × 3.0 × 3.0 mm^3^). This was followed by a field map correction sequence and then T1-weighted images were acquired (repetition time = 7000 ms, echo time = 3500 ms, flip angle = 8°, field of view = 240 mm, slice thickness = 1.0 mm, acquisition matrix = 240 × 240 and voxel size = 1.0 × 1.0 × 1.0 mm^3^). As part of the main ongoing project, diffusion tensor imaging and fast diffusion kurtosis imaging sequences were also acquired with their field map correction. These sequences were not used for this study. Headphones were used to minimize noise exposure and to allow communication, and an eye tracker camera was used to ensure participants remained awake.

### Statistical analysis

Demographic and clinical measures were compared with chi-square tests for categorical variables. The Kruskal-Wallis test was used for ordinal variables; significant between-group differences were followed by pair-wise Mann-Whitney U tests with p < 0.01 per Bonferroni correction for multiple comparisons. For continuous variables, factorial two-way ANOVA was performed if criteria were met, with CD (+/−) and BPD (+/−) as factors. Post-hoc one-way ANOVA with Tukey test was used to assess between-group differences. Analyses were repeated as ANCOVA including as covariates the demographic and comorbidity variables that differed between groups, introducing each in separate models. If after removing outliers and normalizing variables, criteria for ANOVA analysis were not met, we used non-parametric tests. For clinical scales with less than 20% missing values, multiple imputation was performed. If more than 20% of missing values, the subject was eliminated. Analyses were carried out in SPSS 22.0 (SPSS Inc, Chicago, IL, USA).

### Magnetic resonance imaging processing and analysis

T1 images were preprocessed using an in-house pipeline with the software Bpipe (http://cobralab.ca/software/mincbeast_bpipe.html) (31), which uses the MINC Tool-Kit (http://www.bic.mni.mcgill.ca/ServicesSoftware/ServicesSoftwareMincToolKit) and ANTs (32). Briefly, we performed N4 bias field correction (33), linear registration to MNI-space using ANTs, cropped the region around the neck to improve registration quality, followed by transformation back to native space.

The functional images were preprocessed and analyzed using FSL 5.0.8 (34) and AFNI (35). Preprocessing included slice-timing correction, motion correction, field map correction with the FieldMap Topup tool with blip opposite to acquisition direction, brain extraction, segmentation, extraction of the global signal, cerebrospinal fluid signal, white matter signal, physiological noise reduction using aCompCor with 5 PCA factors (36), coregistration, normalization to Montreal Neurological Institute (MNI) stereotactic space, bandpass filtering at 0.01 − 0.08 Hz and smoothing with a 6 mm Gaussian kernel.

We performed seed based analyses, creating 3 mm^3^ spheres using FSL. The seeds for right and left amygdala (lAmy/rAmy) were created at MNI coordinates ±26, 0, 20 (37). The DMN seeds were created using the areas: mPFC, PCC, medial temporal cortex and rostrolateral prefrontal cortex, from the Harvard-Oxford atlas (38). Then we extracted the correlation coefficients between each seed and whole brain using FSL for four regions of interest, which were: mPFC, ACC, right and left insula (rIns/lIns). The second level analysis was done using a two-way ANOVA with CD (+/−) and BPD (+/−) as factors, constrained by five ROIs: bilateral mPFC, ACC and bilateral Insula from Harvard-Oxford atlas. For the main effects and interactions we performed an F-test with FSL randomize (5,000 permutations), followed by pair-wise post-hoc T-tests. For multiple comparison correction we used FWE at 0.05. In the main model we included the following covariates: sex, age, education, current major depressive disorder, current dysthymia and current alcohol use. Finally, as post-hoc analyses, Pearson correlations were performed between correlation scores averaged over clusters with significant FC and scores from the BIS-11 and DERS from each subject.

## Results

### Demographic data

Groups differed significantly in sex and education. Differences in sex possible due to the differences in sample sizes between groups. We also found significant differences in education. These variables were included as covariates for the analyses of ED, impulsivity and FC.

### Clinical findings

The psychiatric comorbidity and medications of the clinical groups are summarized in Supplementary material Table 1S. We found significant differences on current major depressive disorder, current dysthymia, current alcohol use, number of cigarettes consumed per day and antidepressant use. These variables were also included as covariates for analyses of clinical data. In terms of cocaine consumption in the CD+ groups, we found no difference in age of onset (BPD+, CD+ M = 21.0 years, SD = 6.3; BPD-, CD+ M = 22.0, SD 6.18; *U* = 160.5, *p* = 0.563), years consuming (BPD+, CD+ M = 7.4 years, SD = 5.4; BPD-, CD+ M = 8.6, SD 6.18; *U* = 163.5, *p* = 0.624), administration route (smoked: BPD+, CD+ n = 13, 68.4%; BPD-, CD+ n = 11, 57.9%; *X*^*2*^ = 0.11, *p* = 0.737), amount of money spent on cocaine during the last 30 days (BPD+, CD+ M = 149.36 USD, SD = 342.66; BPD-, CD+ M = 185.11, SD 203.90; *U* = 143.0, *p* = 0.180; exchange rate of Mexican pesos to USD at November 7, 2017: 19.14), presence of cocaine in urine (positive: BPD+, CD+ n = 6, 40.0%; BPD-, CD+ n = 7, 53.8%; *X*^*2*^ = 0.537, *p* = 0.724), craving (BPD+, CD+ M = 140.0 points on Cocaine Craving Questionnaire-Now, SD = 37.7; BPD-, CD+ M = 138.0, SD 54.0; *F* = 0.30, *p* = 0.863), and addiction to cocaine severity (BPD+, CD+ M = 33.6 points on the Addiction Severity Index, SD = 15.9; BPD-, CD+ M = 26.3, SD 17.8; *F* = 1.848, *p* = 0.182). The BPD groups did not differ in severity of BPD (Clinical Global Impression-BPD: BPD+, CD+ M = 3.61 points, SD = 1.30; BPD+, CD− M = 3.90, SD 0.92; *F* = 0.394, *p* = 0.535).

For self-reported impulsivity, results of comparison groups are shown in Supplementary Figure 1S. Clinical groups showed lower impulsivity scores compared to controls without psychopathology. In the ANOVA and ANCOVA on BIS-11 total score, the BPD and CD main factors were significant, as was the ANOVA interaction. This interaction shows opposite effects, meaning that the CD factor had lower BIS-11 total scores than BPD. However, the ANCOVA was only significant with some covariates. The best fitting model was obtained with cigarettes/day consumed as a covariate, accounting for 48.9% of the variance. For cognitive and motor impulsivity, the main factor of BPD was significant and the best fit for the model was obtained when cigarettes/day were included, accounting for 41.4% and 36.1% of the variance, respectively. For non-planning impulsivity, the main factor of CD was significant. The analyses are shown in Supplementary Table 2S.

For DERS, the total score did not differ between BPD+CD+ and BPD+CD- groups (BPD+CD+ M = 70.61, BPD+CD- M = 77.67, *U* = 62.0, *p* = 0.348). As indicated in Figure 1, the differences between BPD+CD+ and BPD-CD+, as well as between the BPD-CD+ and BPD+CD- groups approached significance. ANOVA could be performed only for the non-acceptance and goals subscales, after normalizing. For non-acceptance, only the BPD factor was significant (*F = 34.7, p < 0.001*) with an R^2^ of 0.407. With current dysthymia as covariate the R^2^ increases to 0.442 and with cigarettes/day, it diminishes to 0.368 with a nearly significant interaction (*F = 3.78, p = 0.057*), and BPD remaining the only significant effect (*F = 24.31, p < 0.001*). As shown in Figure 1, a negative interaction between BPD and CD factors was found with the goals subscale, meaning counteracting effects of the factors.

**Figure 1.**
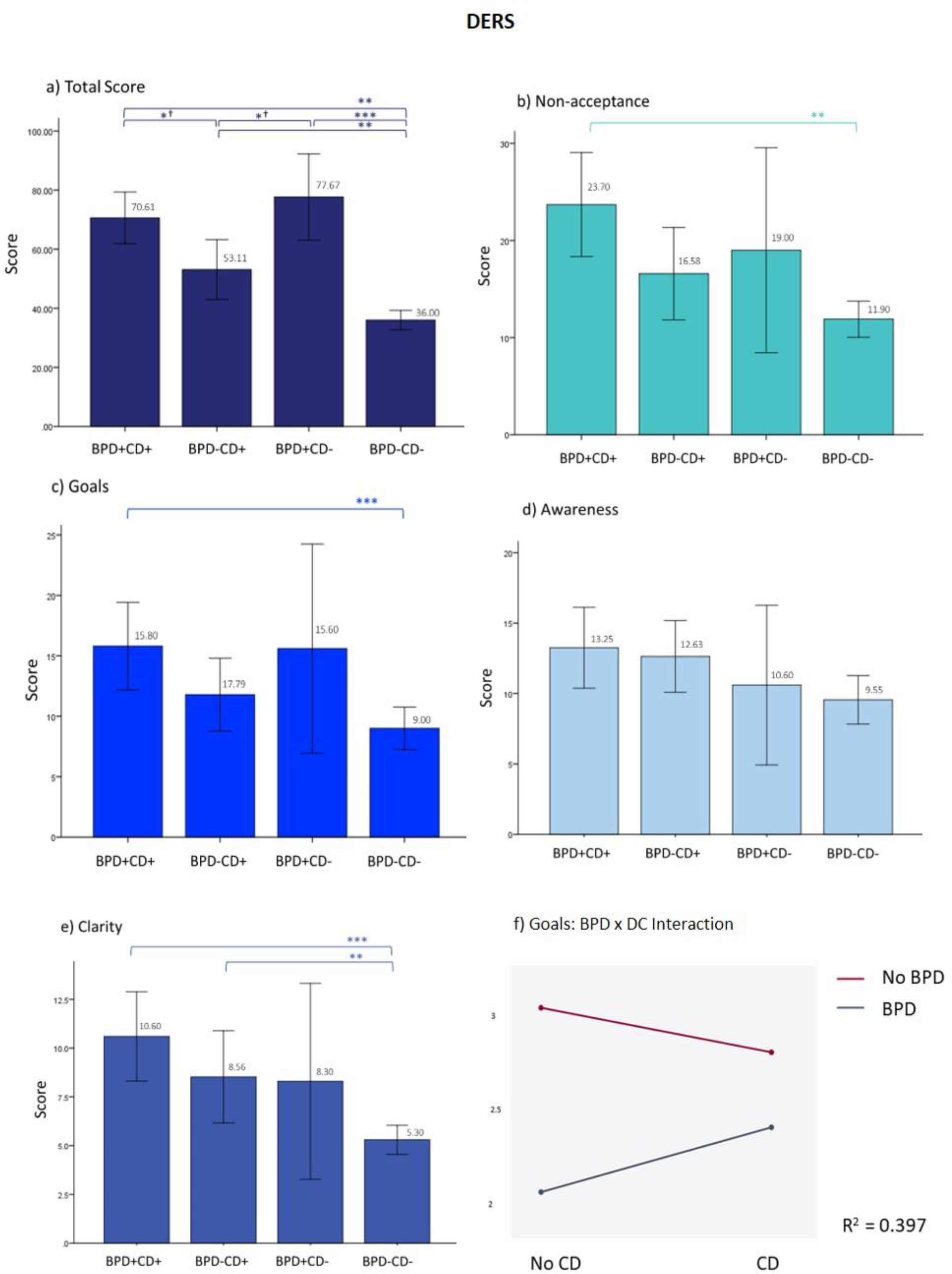
Results from the DERS. a) At total scores, besides the difference between each clinical group and the BPD-CD- group, the difference is near significance between the BPD-CD+ and the BPD+CD+ groups, and the former with the BPD+CD-. b) and c) show a graph with a similar shape than a), but without the significant results. f) Negative significant interaction from the ANOVA at goals subscale (*F* = 6.19, *p* = 0.05) and in the borderline personality disorder factor (*F* = 34.84, *p* < 0.001). When adding cigarettes/day as covariate, the R^2^ improves to 0.421, remaining the interaction significant (*F* = 5.33, *p* < 0.05) and the BPD factor (*F* = 32.04, *p* < 0.001), but not the covariate. On a) to c), *p* value corrected for multiple comparisons to < 0.01. *^†^ p < 0.05, ** p < 0.01, *** p < 0.001. DERS: Difficulties in Emotion Regulation Scale, BPD: borderline personality disorder, CD: Cocaine dependence

### Neuroimaging findings

For the neuroimaging analysis, two participants from the BPD-CD- group were excluded due to poor image quality, leaving n = 67 for analysis. The FC two-way ANOVA analysis showed a significant effect for the interaction between BPD and CD factors in the FC between lAmy and mPFC. This was a negative interaction between the factors, meaning a counteracting effect of each factor: while BPD-CD+ group had an increased FC (significant main effect), BPD+CD- had a decreased FC (near significant main effect). The FC of the BPD+CD+ group was similar to the FC shown by the BPD-CD- group. We also found a significant interaction effect between rAmy and lIns FC, whereby having both factors reduced the FC. Details are shown in Figure 2, Table 2, and Supplementary Figure 2S. The analysis of DMN FC showed no between-group differences for main effects or interactions.

**Figure 2.**
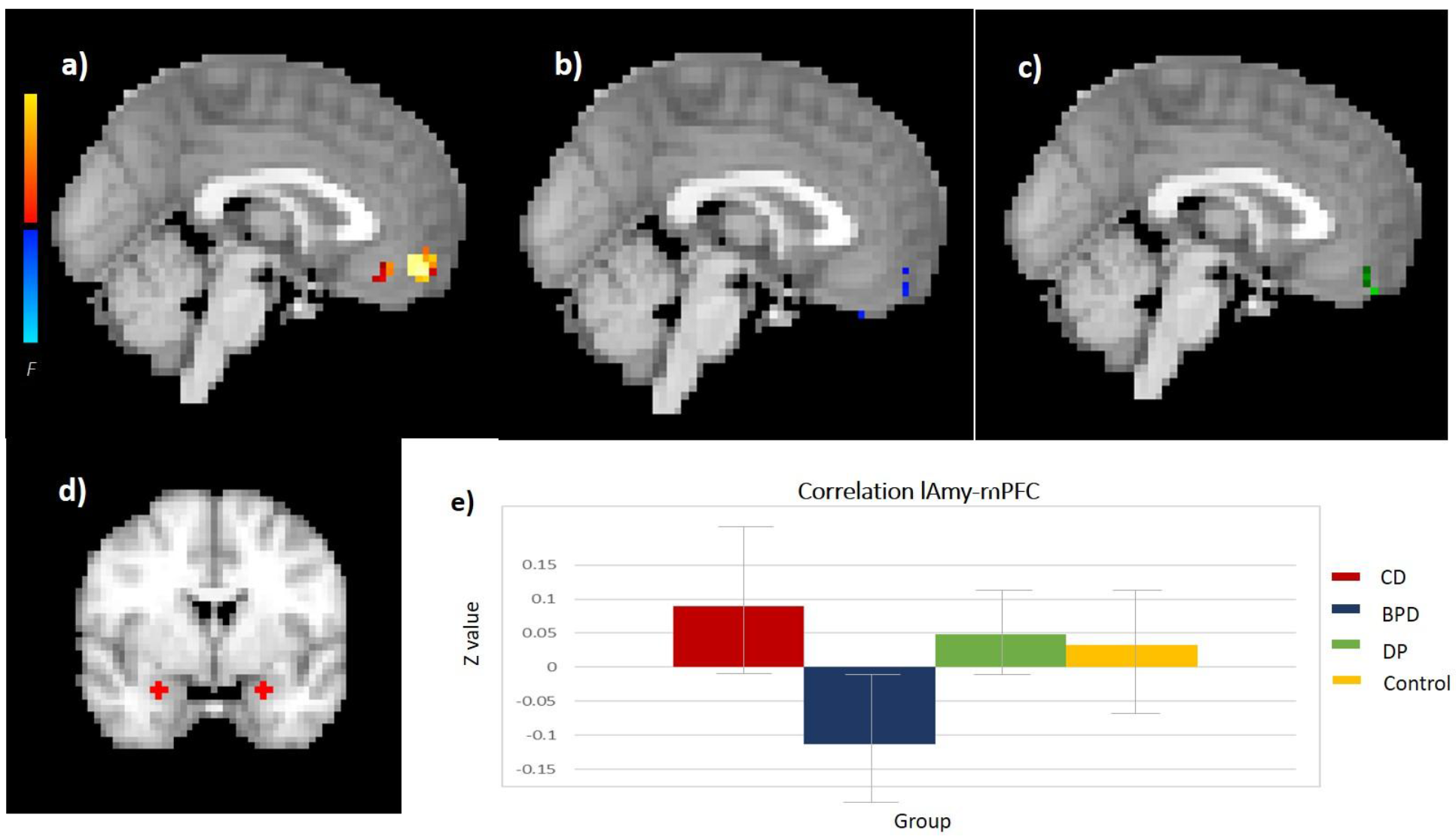
Amygdala connectivity. a-c) Show the significant clusters from lAmy-mPFC connectivity analysis a) the CD effect, b) the BPD effect and c) the interaction. d) Left and right amygdala seeds. e) resting state FC effect sizes for each group with 95% confidence intervals (error bars). lAmy: left amygdala, mPFC: medial prefrontal cortex, CD: cocaine dependence, BPD: borderline personality disorder

**Table 2.** Significant clusters from amygdala connectivity analyses.

| Connectivity analysis<br>Factor | Area | Cluster<br>size<br>(voxels) | Maximum |  |  |  |
| --- | --- | --- | --- | --- | --- | --- |
|  |  |  | Significance (*p*) | MNI coordinates |  |  |
|  |  |  |  | X | Y | Z |
| lAmy – mPFC |  |  |  |  |  |  |
| Interaction | Right frontopolar cortex | 6 | 0.032 | -3 | 48 | -21 |
| Cocaine | Right frontopolar cortex | 42 | 0.008 | 0 | 51 | -15 |
| Borderline<br>personality disorder | Right frontopolar cortex | 1 | 0.055 | -3 | 51 | -9 |
| rAmy-lIns |  |  |  |  |  |  |
| Interaction |  |  |  |  |  |  |
| Cluster 1 | Left insula | 3 | 0.015 | -30 | 9 | -12 |
| Cluster 2 | Left insula | 2 | 0.037 | -36 | -9 | 9 |
| Cluster 3 | Left insula | 1 | 0.049 | -39 | -12 | -12 |

l/r Amy: left/right amygdala, lIns: left insula. For lAmy-mPFC functional connectivity, only one cluster was nearly significant for the borderline factor.

The mean FC obtained from the significant clusters was correlated with BIS-11 and DERS total and subscales scores. We obtained 12 significant correlations shown in Supplementary Table 3S, the strongest being a negative correlation between the cluster from the interaction effect at lAmy-mPFC connectivity with BIS-11 total score. That is, stronger FC between lAmy and mPFC was related to lower scores on self-reported impulsivity. Another negative correlation found was between the first cluster from the interaction effect at rAmy-lIns and non-planning impulsivity. That is, higher rAmy-lIns FC was related to lower non-planning impulsivity. These two correlations are shown in Figure 3.

**Figure 3.**
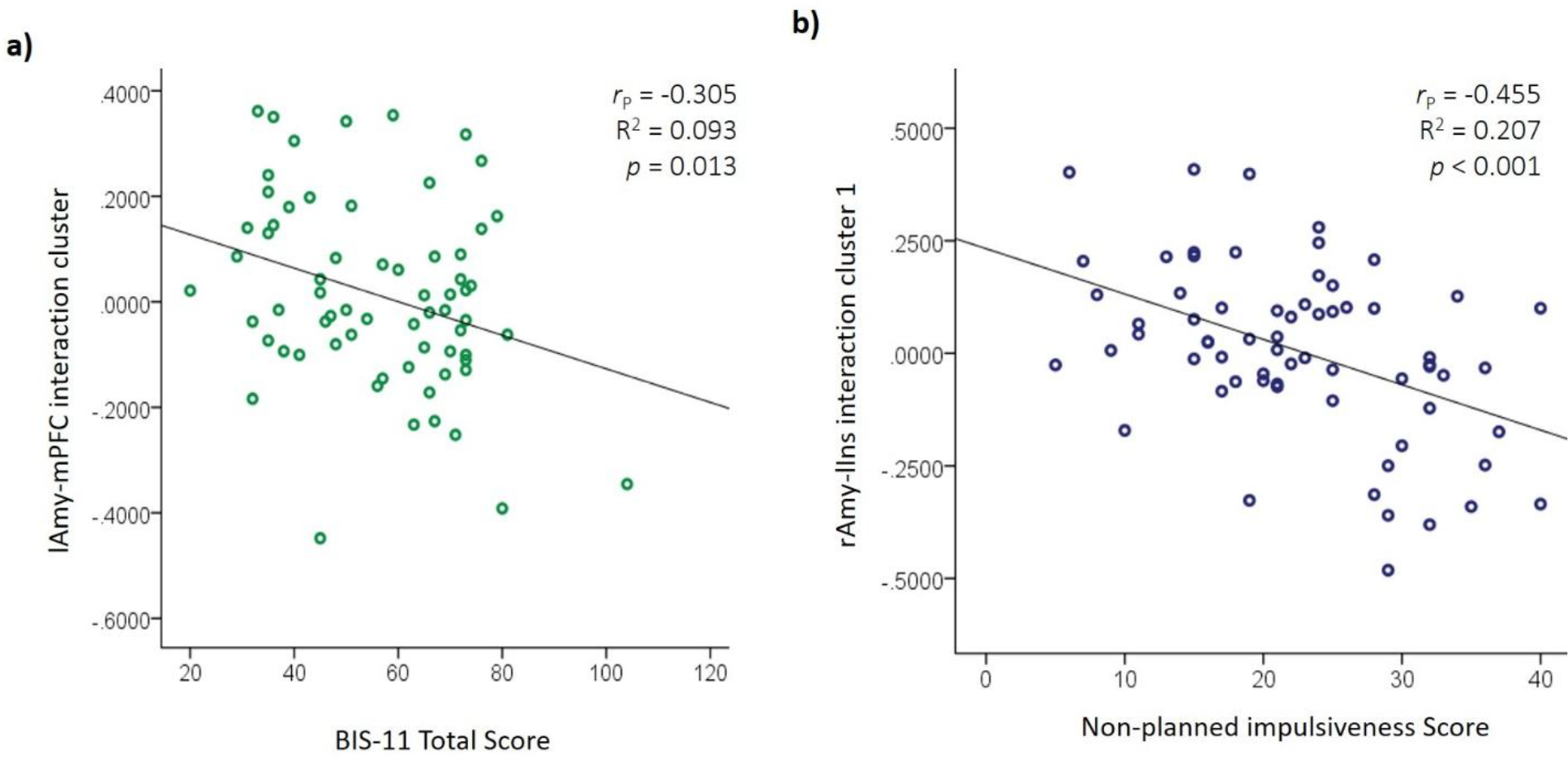
Associations between amygdala functional connectivity and impulsivity. a) lAmy-mPFC connectivity from cluster resulted from the BPD × CD interaction with total BIS-11 score correlation b) rAmy-lIns connectivity from cluster 1 resulted from the BPD × CD interaction with non-planned BIS-11 score correlation. l/rAmy: left/right amygdala, mPFC: medial prefrontal cortex, BPD: borderline personality disorder, CD: cocaine dependence, BIS-11: Barratt Impulsiveness Scale

## Discussion

We sought to understand the psychopathology of BPD with CD, using clinical and FC measures. We found that the BPD+CD+ group resembled the BPD+CD- group in impulsivity and ED scores more than the BPD-CD+ group. We also found that patients with BPD+CD+ displayed a sort of “compensatory” effect in lAmy – mPFC FC, showing a similar FC pattern to the BPD-CD- group, while the rAmy – lIns FC pattern was opposite in the BPD+CD+ group compared to the BPD-CD- group. To our knowledge, this is the first study to investigate the effect of this DP using clinical and neuroimaging measures.

In terms of psychiatric comorbidities, we found greater prevalence of current major depressive episode and dysthymia in the BPD+CD- group. This could reflect the greater proportion of women in that group, as mood disorders are more frequent among women than men with BPD (39–42). We also found higher current alcohol consumption in the BPD-CD+ group. As reported, the presence of a substance use disorder increases the likelihood of abusing other substances, especially for males (1, 40); our BPD-CD+ group had the highest proportion of males. The number of cigarettes consumed per day was highest in the BPD+CD+ group. Nicotine consumption has been associated with use of other substances and with BPD (43). Introducing these variables as covariates in the remaining analyses can reduce their influence in the inference models, as they are difficult to control in composition of the groups due to the nature of the study population.

Impulsivity and ED are core characteristics of BPD. Impulsivity was described as the main predictor of borderline symptomatology in a seven year follow-up (15), while ED predicts aggressive behavior even more than impulsivity (44). With respect to CD, impulsivity predicts higher cocaine consumption (14) and increases the risk for a stimulant use disorder (13); during brief periods of abstinence, patients with CD show ED (17). In previous studies of BPD and CD, the DP group presented more severe difficulties with sexual risk behaviors (9) and greater attentional bias toward cocaine cues under an emotional stress condition (10) than groups with only one of the two disorders. Taking this into account, clinically we expected an additive effect of BPD combined with CD on impulsivity and ED, however, we found similar scores in both constructs for patients with BPD+CD+ and BPD+CD-. This suggests either that BPD is their main pathology or that CD may be compensatory, although these hypotheses are clearly speculative.

In terms of neuroimaging, we found a significant interaction in lAmy – mPFC FC with the BPD and CD factors producing opposite effects and reducing the FC in the BPD+CD+ group to the level of the BPD-CD- group. The FC strength of this circuit was negatively correlated with impulsivity and ED. These findings make sense in light of what has been described in literature about amygdala – mPFC FC in relation to emotion regulation and impulsivity and with our clinical findings. Amygdala – mPFC connectivity is important for top-down emotion regulation (45, 46), a process that is impaired in patients with BPD (47) and in CD (17). Functional connectivity between these brain areas has been linked to impulsive aggression and with MRI signal variance modified by sex (48). The functional connection between these structures has also been suggested to facilitate the urgency to consume the substance of interest during abstinence in individuals with dependence, sending to a so-called “impulsive system”, consisting of the amygdala and nucleus accumbens, signals that magnify the value of somatic marker representations (49). In the BPD+CD- group, our results agree with previous studies that did not find differences in amygdala – mPFC FC compared with BPD-CD- (50, 51). Amigdala – mPFC FC has been reported to be lower in CD compared to controls (20), while we found it non-significantly higher. However, their sample was older and without any psychiatric comorbidity, and their resting state fMRI preprocessing also differed from ours.

Another relevant finding was the significant interaction effect in rAmy – lIns FC, where having DP significantly diminished their FC, in an additive manner. This FC was negatively correlated with ED and impulsivity scores, especially with non-planned impulsivity, which relates to self-control and cognitive complexity (52). The insula has been linked to emotion regulation (53) and impulsivity (54, 55), and is a key component of the salience network which is activated in sensory stimulus-guided goal-directed behaviors (56). In BPD patients, insula function has been related to emotional processing of pain (57). In substance use disorders, craving involves a dysregulation of afferent projections from the insula to amygdala and related structures (58). Our finding of additive effects of both factors in DP patients and the correlations with clinical measures might help explain the greater attentional bias to cocaine-related visual stimuli under emotional stress found in another sample (10). Under this model, aberrant rAmy - lIns FC would impair emotion regulation under stress and make cocaine cues more salient.

Overall, the resting state FC results fit with our clinical findings, with the notion that CD may be an empirical method of self-regulating Amy - mPFC brain connectivity in BPD, to the detriment of other cortical connections such as rAmy - lIns, and therefore, enhancing attentional bias to cocaine cues in DP patients. Clinical guidelines and the literature indicate that when a DP diagnosis is present, both disorders must be treated in an integrated manner (59, 60). However, mental health teams able to manage dual diagnosis patients remain scarce, so that currently the common practice is to treat the substance use disorder predominantly or exclusively, even in specialized clinics (61).

### Limitations

The main limitation of our study is the small sample size of groups, especially the BPD+CD- group, which limited our statistical power. When we introduced covariates, a number of significant results changed, making it difficult to assess their impact. Despite this, our sample was uniform on illnesses severity and on most clinical variables. In neuroimaging, sample sizes of at least 22 subjects per group are recommended for task-based studies (62). No such estimates are available for resting state FC studies, and reaching those sample sizes would have been extremely difficult given the types of patients we studied. Therefore, we limited our analyses to candidate-circuit analyses based on prior evidence, greatly reducing our multiple-comparisons problem.

Another issue is that most CD studies have included only males, and BPD studies have tended to include only females, while we included both sexes. However, we were still not able to analyze sex as an independent factor, as suggested in previous research (9, 10, 48). A further limitation is that our participants were recruited from a clinical population seeking treatment in specialized units, making it difficult to generalize the results to the broader population of patients not in treatment. This, however, is not unique to our study. We used only self-reported measures for ED and impulsivity. However, a trained psychiatrist who knew the patients reviewed their responses. Despite these limitations, this is the first study that examines impulsivity, ED and FC in the dual pathology of CD and BPD.

In summary, we found that patients with dual pathology showed similar difficulties in impulsivity and ED as those with BPD without CD, and they had similar resting state amygdala – mPFC FC to healthy controls, while the dual pathology group had reduced amygdala – insula FC than all other groups. This suggests the speculative hypothesis that cocaine consumption may be a form of self-medication in BPD to normalize amygdala – mPFC FC, but that it also affects other brain connections like the amygdala – insula circuit. If this preliminary observation is replicated, it would suggest that treatment approaches to this dual pathology need substantial revision.

## Acknowledgements

The study was supported by grants from the National Council of Science and Technology of Mexico CONACYT-FOSISS project No. 0201493 and CONACYT-Cátedras project No. 2358948. We also thank the support of Francisco Xavier Castellanos, M.D. for editorial assistance and Ms. Ana Teresa Martínez Alanís. Recruitment of patients was supported by the Toxicologic Medical Unit (treatment clinic) Xochimilco in Mexico City and the National Institute of Psychiatry “Ramón de la Fuente Muñiz”.

